# BonA from *Acinetobacter baumannii* forms a divisome-localized decamer that supports outer envelope function

**DOI:** 10.1101/2020.09.01.278697

**Authors:** Rhys Grinter, Faye C. Morris, Rhys A. Dunstan, Pok Man Leung, Matthew Belousoff, Sachith D. Gunasinghe, Simone Beckham, Anton Y. Peleg, Chris Greening, Jian Li, Eva Heinz, Trevor Lithgow

## Abstract

*Acinetobacter baumannii* is a high-risk pathogen due to the rapid global spread of multi-drug resistant lineages. Its phylogenetic divergence from other ESKAPE pathogens means that determinants of its antimicrobial resistance can be difficult to extrapolate from other widely studied bacteria. A recent study showed that *A. baumannii* upregulates production of an outer-membrane lipoprotein, which we designate BonA, in response to challenge with polymyxins. Here we show that BonA has limited sequence similarity and distinct structural features compared to lipoproteins from other bacterial species. Analyses through X-ray crystallography, small-angle X-ray scattering, electron microscopy, and multiangle light scattering demonstrate that BonA has a dual BON-domain architecture and forms a decamer via an unusual oligomerization mechanism. This analysis also indicates this decamer is transient, suggesting dynamic oligomerization plays a role in BonA function. Antisera recognizing BonA shows it is an outer membrane protein localized to the divisome. Loss of BonA modulates the density of the outer membrane, consistent with a change in its structure or link to the peptidoglycan, and prevents motility in a clinical strain (ATCC 17978). Consistent with these findings, the dimensions of the BonA decamer are sufficient to permeate the peptidoglycan layer, with the potential to form a membrane-spanning complex during cell division.

## Introduction

*Acinetobacter baumannii* is a notorious ‘red alert’ pathogen, considered an urgent threat to human health by international infectious disease control agencies [1-4]. As a member of the gammaproteobacterial family *Moraxellaceae, A. baumannii* is genetically and physiologically divergent from well-studied model Gram-negative *Enterobacteriaceae* such as *Escherichia coli. A. baumannii* has a unique cell envelope that allows it to survive exposure to disinfectants and desiccation that readily kill other bacterial species, allowing it to persist for long periods on artificial surfaces in hospitals [5, 6]. Additionally, *A. baumannii* is notorious for its innate and acquired antibiotic resistance [2]. It is currently estimated that as many as 50% of all *A. baumannii* infections in the USA are caused by strains resistant to carbapenems and many strains acquire polymyxin resistance during treatment [7, 8].

Like other Gram-negative bacteria, *A. baumannii* has a cell envelope consisting of an inner and outer membrane. This dual membrane encloses the periplasm, a crowded compartment that contains a thin layer of peptidoglycan [9]. The outer membrane of *A. baumannii* is an intricate structure, consisting of an asymmetric lipid bilayer with an inner leaflet composed of phospholipids and an outer leaflet composed of lipooligosaccharide (LOS) [10]. The LOS derived surface of the outer membrane acts as a barrier to hydrophobic molecules [11]. In addition to LOS and phospholipids, the outer membrane contains numerous proteins that are either integrated into or anchored onto the membrane [12].

To maintain the integrity of the outer membrane, Gram-negative bacteria actively maintain its lipid asymmetry and coordinate its biogenesis rate with the overall rate of cell growth. Additionally, the outer membrane must be constricted in conjunction with the peptidoglycan cell wall during division [13]. To achieve this, Gram-negative bacteria have evolved a network of interlinked pathways for the construction and maintenance of the outer membrane [12, 14-22]. Despite considerable progress in understanding how these pathways function in *E. coli*, in many cases, the proteins that constitute them are not well characterized, and additional pathways likely remain to be identified [12, 20, 23]. In species divergent from *E. coli*, such as *A. baumannii*, these knowledge gaps are much more substantial.

Among these knowledge gaps is the role of dual-BON domain proteins, a widespread family of outer envelope proteins in Gram-negative bacteria. Dual-BON family proteins contain a pair of <u>B</u>acterial <u>O</u>smY and <u>N</u>odulation (BON) domains, which fold into a conserved α/β sandwich [24]. They possess a signal peptide targeting them to the periplasm, and some family members possess a lipobox with an N-terminal acylated cysteine, mediating peripheral outer membrane association [25, 26]. They lack conserved residues indicative of an enzyme active site, though some family members bind phospholipids [27, 28]. Archetypical members of this dual-BON domain family are the outer membrane-associated lipid-binding protein DolP (formerly YraP) and the soluble periplasmic protein OsmY, both of which play a role in the construction and maintenance of the bacterial outer envelope [25, 26, 29]. OsmY is an abundant periplasmic protein in *E. coli* induced in response to stressors such as osmotic shock, heat shock, acidic pH, and bile salts [25, 30]. Recently, it was shown that OsmY functions as a chaperone, enhancing the stability of periplasmic proteins and the assembly of a subset of outer membrane proteins [31].

DolP is a lipoprotein widely present in Gram-negative bacteria. In *E. coli* and *Neisseria meningitidis*, it localizes to the inner leaflet of the outer membrane via an N-terminal lipid anchor [32-34]. DolP was initially identified in *E. coli* as a lipoprotein whose expression is induced under cell envelope stress and it forms part of the σ^E^ regulon [35]. Mutants of *E. coli, N. meningitidis*, and *Salmonella enterica* lacking DolP are compromised in outer membrane integrity, rendering the cells more sensitive to agents like the detergent SDS or the antibiotic vancomycin [26, 28, 33, 35, 36]. Likely as a result of impaired outer membrane integrity, loss of DolP leads to attenuation of virulence in rodent models of infection [26]. Despite the phenotypic characterization of mutants lacking DolP, suggesting a role in outer membrane biogenesis [37, 38], the specific biochemical function of DolP remains to be established. In *E. coli*, DolP is recruited to the site of cell division [32]. This recruitment is required for the regulation of cell wall remodeling during cell division [32, 39]. A recent study resolved the structure of DolP from *E. coli*, showing that it conforms to a dual-BON domain architecture and is monomeric [28]. This study demonstrated that DolP binds anionic phospholipids via α-helix 1 of its C-terminal BON domain, and that phospholipid binding is required for its function and localization. Interestingly, despite relatively low overall sequence identity, the sequence of this lipid binding helix is highly conserved in DolP from *N. meningitidis [29]*, suggesting a conserved function for these proteins.

The focus of this study is a dual-BON domain protein synthesized by *A. baumannii*. Our previous work has shown that this bacterium can become resistant to the LOS-binding antibiotic polymyxin through mutations that prevent LOS production [40]. These mutants survive with an outer membrane where phospholipids compose the only lipid species in both leaflets of the membrane [40]. In both wild-type polymyxin treated cells and in polymyxin resistant LOS-deficient mutants, a dual-BON domain family lipoprotein protein (HMPREF0010_02957, ABBFA_002498) is upregulated [41, 42]. This suggests that this protein, which we designate BonA, plays a role in adapting to the effects of polymyxin on the *A. baumannii* outer envelope, and to the loss of LOS. BonA is only distantly related to either DolP or OsmY and we show that, unlike DolP, its loss does not lead to a gross outer membrane permeability defect. Alternatively, *A. baumannii* mutants lacking BonA have an altered outer membrane structure and a defect in cell motility. Like DolP, single-cell imaging of *A. baumannii* indicates that BonA is localized to the divisome. However, BonA lacks conserved amino acids that mediate phospholipid binding by DolP, indicating a divergent function at this location. Through structural and biophysical analysis, we show that BonA forms a decamer and that this oligomerization is stabilized by a novel mechanism, involving rearrangement of the BON-domain fold. This oligomerization provides a rationale for explaining how BonA functions in the absence of a conserved lipid-binding motif or active site and is consistent with a scaffold or chaperone function for the protein. Based on its unique structure, dynamic oligomerization, and role in outer membrane maintenance, this study establishes BonA as a third branch of the dual-BON domain family, distinct from OsmY and DolP found in other bacteria.

## Results

### BonA from *A. baumannii* is a member of a diverse family of dual-BON domain outer-membrane lipoproteins

Analysis of *A. baumannii* genomes showed that they encode only one BON domain family protein. The amino acid sequence of this lipoprotein contains dual-BON domains, a terminal lipobox with an acyl-anchoring cysteine residue, and N- and C-terminal extensions (Figure 1A). BonAshows high sequence divergence from DolP from *E. coli* (23% identity) and *Neisseria* spp. (24% identity), and is even more distantly related to OsmY from *E. coli* (20% identity) (Table S1). A phylogenetic tree confirmed the distant evolutionary relationship between BonA and other dual-BON domain lipoproteins identified in a HMMER search of the reference proteome database (Figure 1B, Table S2) [43]. BonA belongs to a distinct clade clustering with proteins from other members of the family *Moraxellaceae*. A C-terminal proline-rich extension is present in BonA and other related sequences from *Acinetobacter* and *Moraxella* species but is absent from DolP and OsmY (Figure 1A, Figure S1).

**Figure 1.**
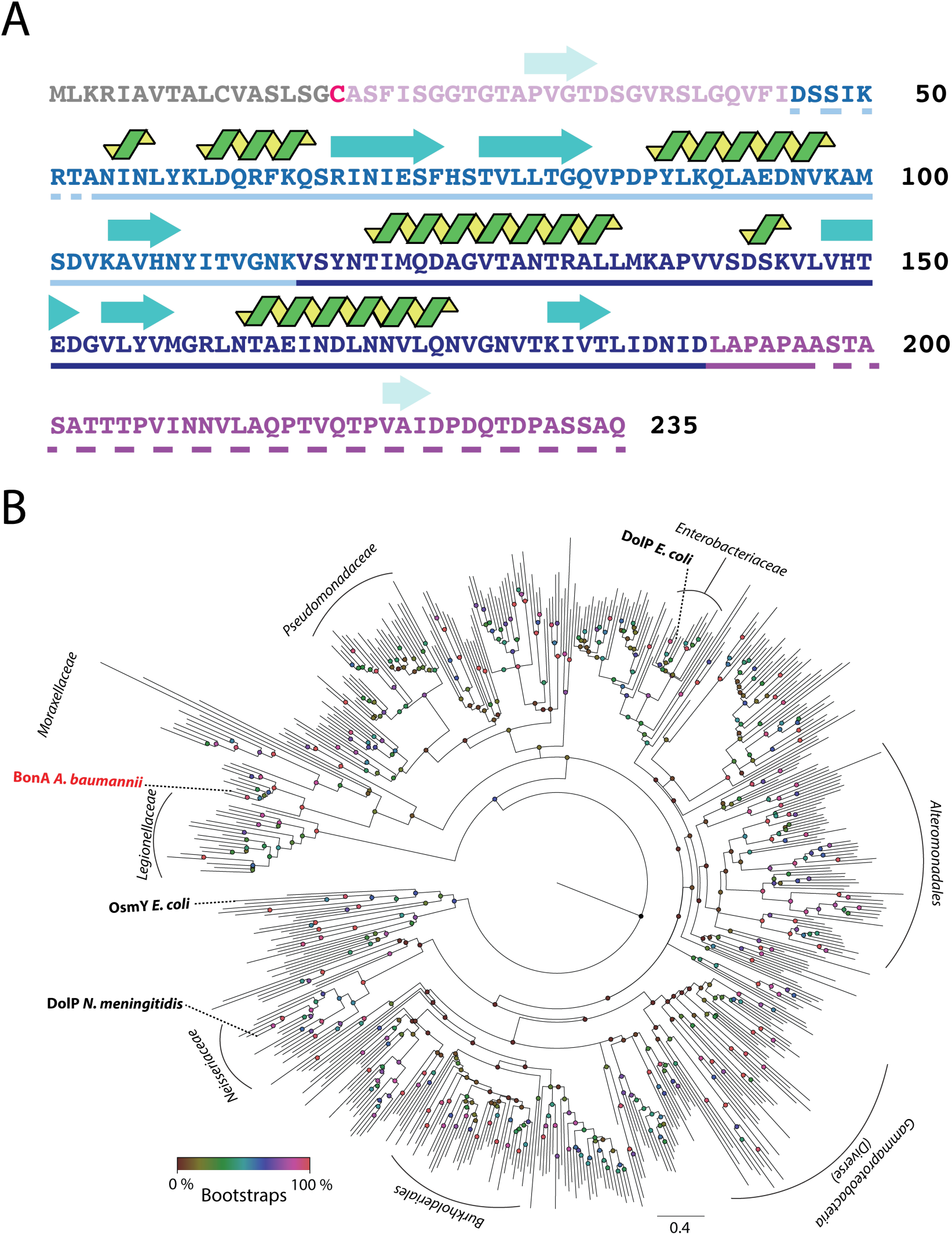
The sequence, secondary structure and molecular phylogeny of BonA. (A) The amino acid sequence of BonA showing secondary structure (β-sheet = blue arrows, α-helix = green spirals; predicted or based on the BonA-27N crystal structure), the location of BON1 (light blue) and BON2 (dark blue), regions largely lacking predicted structure (purple) and the signal peptide (grey) and acyl-anchored cysteine (red). Amino acids present in BonA-27N are underlined, solid for those resolved in the crystal structure, and dashed for disordered regions. (B) A maximum-likelihood phylogenetic tree of BonA homologs, shown in Table S1, showing the relatedness of BonA to the characterized family member DolP from *E. coli* and *N. meningitidis*. The clade containing the distinct dual-BON domain family member OsmY from *E. coli* was used to root the tree. Nodes are color-coded according to bootstrap values based on 100 replicates.

### BonA is localized to the divisome and its deletion prevents motility

The distant evolutionary relationship between BonA and other dual-BON family proteins poses the question of whether these proteins share a conserved function. To address this we sought to determine the subcellular localization and physiological role of BonA. Mutants of the well-characterized *A. baumannii* type strain ATCC 19606 and clinical isolate ATCC 17978 were constructed (Δ*bonA*). Antibodies raised to BonA detected the protein in wildtype *A. baumannii* ATCC 19606, but not in the Δ*bonA* strain when membrane extracts were analyzed by SDS-PAGE and immunoblotting (Figure 2A). To monitor the subcellular localization of BonA, cell membrane extracts were fractionated via a sucrose gradient, followed by immunoblotting, revealing that BonA is localized to the outer membrane as predicted by its N-terminal lipobox (Figure 2B).

**Figure 2.**
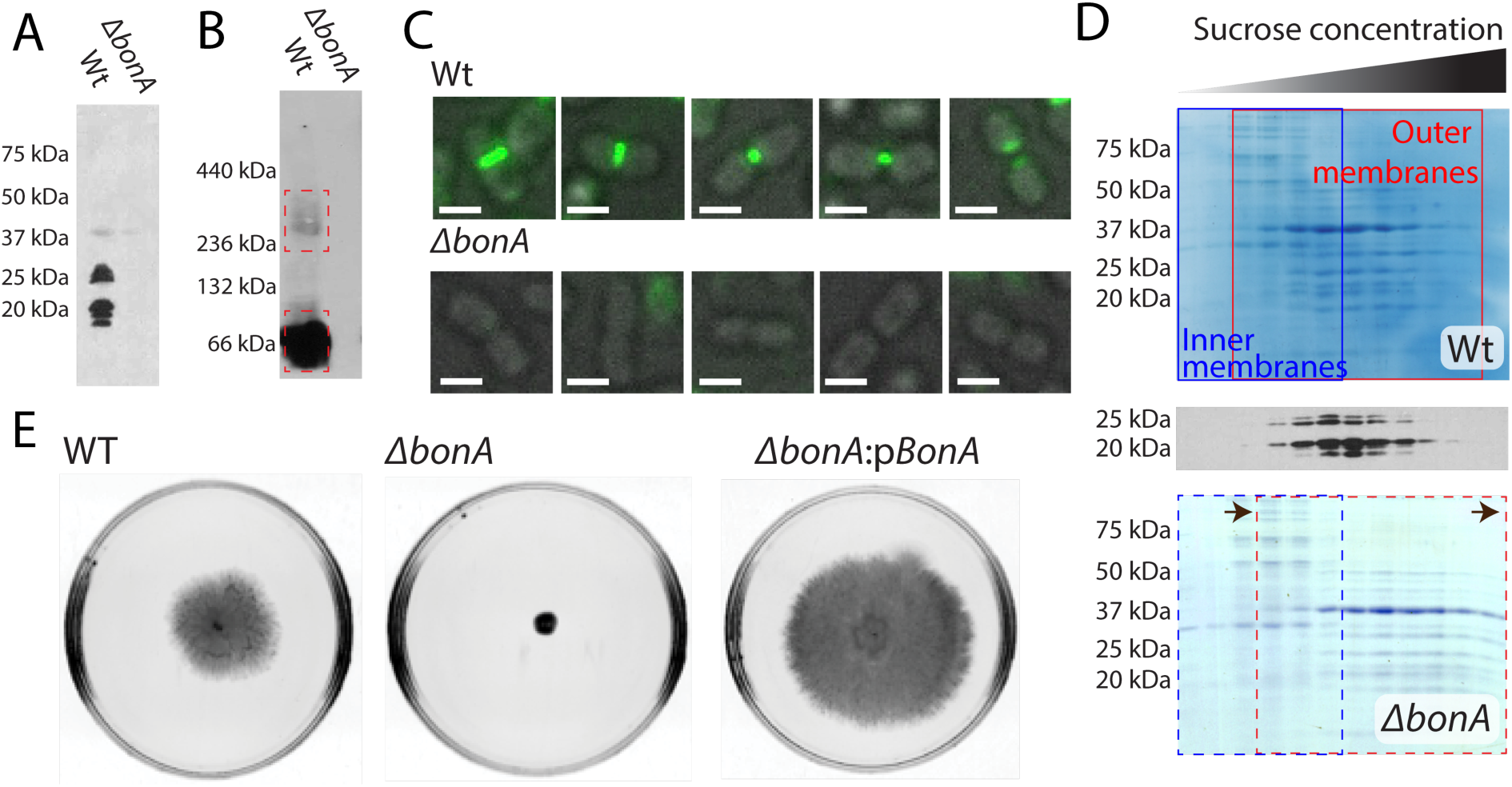
The cellular localization of BonA and phenotypes associated with loss of BonA in *A. baumannii*. (A) An SDS-PAGE Western blot of total cellular membranes from wildtype and *ΔbonA A. baumannii* ATCC19606 with an anti-BonA antibody, showing BonA is membrane localized. (B) A blue-native PAGE Western blot of membranes as in panel A, showing BonA adopts a dimer and higher MW species when purified from native membranes. (C) Immunofluorescence microscopy of wildtype and *ΔbonA A. baumannii* ATCC19606 using an anti-BonA antibody, showing BonA is localized to the site of cell division; Scale bar = 2 μM. (D) Sucrose gradient separation of membranes from Panel A/B showing that BonA is associated with fractions containing the outer membrane and that in the *ΔbonA* the outer membranes exhibit a higher density on the sucrose gradient. (E) Semi-solid motility assay plates of *A. baumannii* ATCC17978, showing that the *ΔbonA* is non-motile compared to the wildtype and complemented mutant, where expression of BonA from pWH1266 restores this phenotype.

While the relative abundance of proteins present in *A. baumannii* ATCC 19606 Δ*bonA* membranes was similar to wildtype, the outer-membrane fraction from the Δ*bonA* strain progressed markedly further into the sucrose gradient. This suggests that its structure or composition is altered, leading to an increase in density (Figure 2B). However, no significant increase in sensitivity to SDS, vancomycin, or tetracycline was observed (Table S3), suggesting that loss of BonA does not impair the integrity of the outer membrane in *A. baumannii*. Loss of motility on a swarm plate assay was observed in the *A. baumannii* ATCC 17978 *ΔbonA* strain, which could be complemented by the addition of *bonA* in trans (Figure 2E). ATCC 19606 is non-motile in this assay, and so this phenotype could not be tested in this strain (Figure S2). While *A. baumannii* lacks flagella, twitching motility is observed in some strains of this species, thought to be mediated by the type IV pilus [44]. Type IV pili are dynamic protein filaments that are assembled and secreted from the cell via a large protein complex that spans the bacterial cell envelope [45]. The loss of twitching motility observed in the ATCC 17978 *ΔbonA* mutant suggests that BonA plays either a direct or indirect role in the assembly or function of this molecular machine.

Like BonA, DolP is a lipoprotein anchored to the outer membrane. DolP is localized to the divisome where it plays a role in regulating peptidoglycan remodeling during cell division [32]. To determine if BonA shares a common localization, we used the antibodies to monitor BonA in single cells via immunofluorescence microscopy. Consistent with localization to the divisome, fluorescence corresponding to BonA was observed as a central band in what appeared to be elongated, early-stage dividing cells. This band constricted in concert with the cell-division septum (Figure 2C). No fluorescence beyond background was observed in *ΔbonA* cells (Figure 2C). To investigate the native structure of BonA, membrane extracts were solubilized in detergent and analyzed by blue-native PAGE. The vast majority of BonA was detected at a molecular weight of ∼60 kDa, consistent with a dimer or trimer of the 23 kDa protein (Figure 2D). Longer exposure of the immunoblots revealed a smaller proportion of BonA was detected as a larger oligomeric species (250-300 kDa).

### The crystal structure of BonA indicates functional divergence from other dual-BON proteins

To gain insight into the structural organization of BonA compared to other dual-BON proteins, as well as its architecture at the outer membrane, we solved its crystal structure. Crystal trials were performed with full-length BonA as well as several truncation constructs. High-quality crystals were only obtained for N-terminally truncated BonA, missing the 27 amino acids after its lipid anchoring cystine. The structure of this protein, designated BonA-27N, was solved at 1.65 Å by single-wavelength anomalous dispersion (SAD) phasing, using selenomethionine labeled protein. The structure of BonA-27N was built and refined from the resulting electron density maps (Table S4, Figure 3A). The crystal structure of BonA-27N consists of two α/β-sandwich BON domains that interact extensively via the external face of their three-strand β-sheets (Figure 3A). In contrast to the structure of DolP in which both domains adopt a canonical BON domain fold [28], in the BonA structure, α-helix 1 (αH1) of BON domain 1 (BON1) does not adopt the expected BON domain conformation of running parallel to the BON domain β-sheet. Rather it is displaced from the rest of the domain (Figure 3A). The 39 amino acids of the C-terminal extension of BonA-27N (AAs 196 to 235) are disordered in the crystal structure. This region of BonA is not present in DolP or OsmY and is predicted to be largely unstructured (Figure 1A).

**Figure 3.**
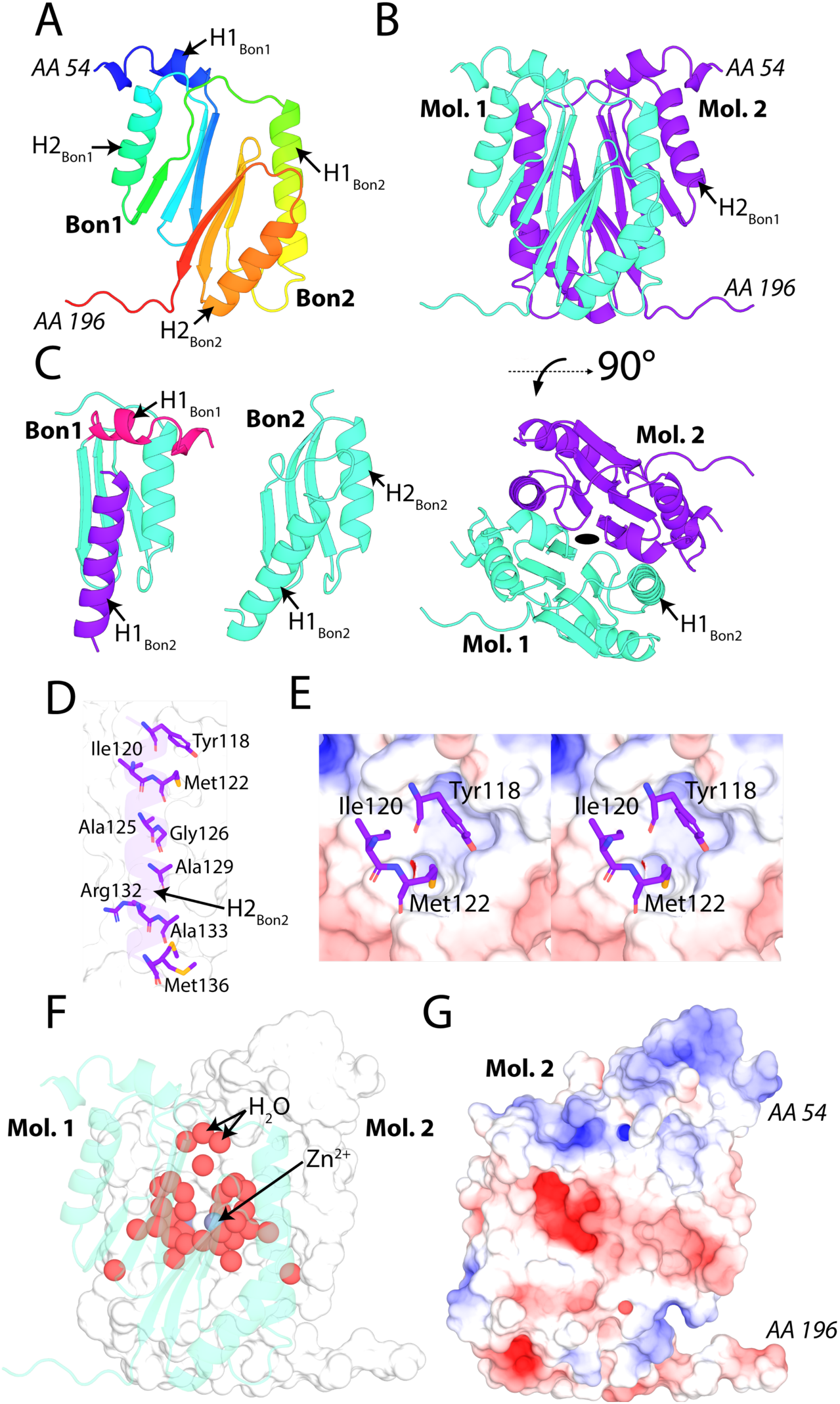
The crystal structure of BonA-27N reveals a dual-BON domain architecture that dimerizes via an α-helix swap mechanism. (A) The crystal structure of BonA-27N shown as a rainbow cartoon N-terminus (blue) to C-terminus (red) displays a dual BON domain architecture with displaced α-helix 1 (αH1) of BON domain 1. (B) The dimer of BonA-27N observed *in crystallo*. (C) A key interface of the BonA-27N dimer involves the displacement of αH1 of BON1 by α-helix 1 (αH1) of BON2 of the opposing BonA molecule. αH1 of BON2 is amphipathic and interacts with the opposing molecule largely through hydrophobic interactions shown in (D) and (E). As shown in (F) and (G) the BonA dimer interface is highly hydrated and consists of both hydrophobic and polar interactions.

Analysis of BonA-27N crystallographic symmetry reveals that it exists as a dimer, aligned with the crystallographic two-fold axis (Figure 3B). Analysis with the molecular interface prediction tool PISA [46] predicts that this interface is *bona fide* (Table S5). The symmetrical BonA-27N dimer interacts via an extensive interface encompassing both BON domains (Figure 3B). The interface is stabilized by α-helix 1 (αH1) of BON domain 2 (BON2), which substitutes for the displaced αH1 of BON1, thus completing the α/β-sandwich fold of BON1 (Figure 3C). This interaction of αH1 of BON2 with BON1 is largely mediated by hydrophobic interactions (Figure 3D), with Tyr118 and Met122 of αH1 of BON2 extending deeply into a hydrophobic pocket created by the displacement of αH1 of BON1 (Figure 3E). While the interactions between αH1 of BON2 and BON1 are largely hydrophobic, the dimer interface of BonA-27N is mediated by a mixture of interaction types, including 14 hydrogen bonds and four salt bridges (Figure 3G, Table S5). The interface also contains two symmetrical, highly solvated pockets, which trap a total of 34 water molecules, as well as two Zn^2+^ ions which were present at a high concentration in the crystallization solution (Figure 3F).

Altogether, these findings show that BonA is structurally and functionally distinct from other dual-BON family proteins. In contrast to BonA, the structure of DolP from *E. coli* reveals it is monomeric and its BON1 domain adopts the canonical α/β-sandwich fold. Additionally, rather than mediating oligomerization, αH1 of BON2 of DolP from *E*.*coli* is responsible for binding to anionic phospholipids present in the outer-membrane, via residues that are not conserved in BonA [28]. These differences indicate BonA is structural and functionally divergent to the DolP branch of the dual-BON domain protein family. A recent study by Wu et. al. supports the physiological relevance of the BonA dimer, demonstrating via a global proteomic approach that intermolecular interaction occurs between BonA molecules in *A. baumannii* cells [47]. This study identified intermolecular crosslinks between lysines 50, 59, and 65 of neighboring BonA molecules in *A. baumannii* cells. In the BonA-27N structure, lysine 59 and 65 are located in αH1 of BON1 and are within proximity to their dimer equivalent in our BonA-27N structure (Figure S3). Lysine 50 is unresolved in the crystal structure, but given this region of BonA is crucial for oligomerization, it is also a plausible candidate for crosslinking based on our data.

### BonA decamerises under physiological conditions through interactions mediated by its N-terminal extension

Our structural and biochemical analysis indicated that BonA oligomerizes in *A. baumannii* cells and as a recombinant protein. To investigate the oligomeric state of BonA, the mature recombinant protein (lacking its signal sequence) was analyzed by size exclusion chromatography (SEC). In the absence of detergent, BonA migrated predominately as a high-molecular weight species, with some disassociation to a lower-molecular weight species observed. To gain a more precise understanding of the molecular weight of this oligomer, purified BonA was analyzed by analytical ultracentrifugation, revealing the presence of a single species with a molecular mass of approximately 240 kDa (Figure S4A). The molecular mass of the BonAoligomer was confirmed by size-exclusion chromatography coupled multiangle laser-light scattering (SEC-MALS), which indicated this species has a molecular mass of 233 kDa, which is consistent with a decamer, while the smaller species has a mass of 23 kDa, corresponding to a BonA monomer (Figure 4B). To determine if the N or C-terminal extensions flanking the core BonA BON domains were responsible for oligomerization, truncation constructs lacking the N-terminal 27 amino acids succeeding the lipobox and/or the C-terminal 45 amino acids were analyzed via SEC-MALS (Figure 1A, Figure 4A). Removal of the C-terminal extension increased the tendency of BonA to aggregate but did not affect the oligomeric state of the protein, with a decamer of 205 kDa observed for the truncated protein (Figure 4C). Conversely, loss of the 27 N-terminal amino acids abrogated oligomerization, with only a monomeric species of ∼22 kDa observed (Figure 4D). Loss of both the N- and C-terminal regions also resulted in a monomeric protein, further confirming the role of the N-terminus of BonA in oligomerization (Figure 4E). In conclusion, BonA forms a decamer that requires its N-terminal extension and undergoes spontaneous disassociation into a monomeric species in solution. The monomeric nature of BonA-27N in solution contrasts with the dimer observed in its crystal structure, suggesting that weak interactions between monomers of this truncated protein are selected for during crystallization.

**Figure 4.**
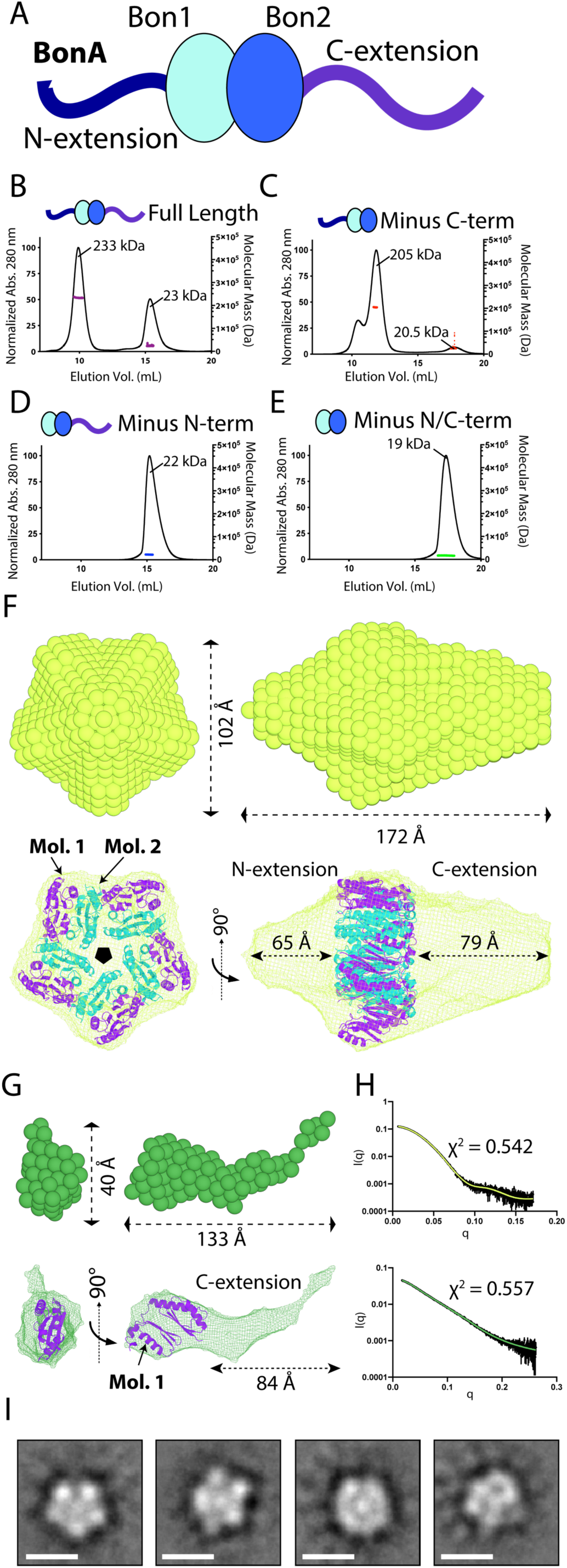
BonA forms a decamer with pentameric symmetry mediated by its N-terminus. (A) A cartoon schematic of BonA showing its two central BON domains, with N and C-terminal extensions with limited predicted secondary structure. SEC-MALS experiments showing that (B) full-length soluble BonA and (C) a 45 amino acid C-terminally truncated variant are predominantly decamers, with some disassociation into a monomer. Conversely, (D) 27 amino acid N-terminally truncated and (E) 45 amino acid C and 27 amino acid N-terminally truncated variants both are monomers. (F) A bead model of the full-length BonA decamer modeled from SAXS data with C5 symmetry imposed (top), and a mesh representation of this bead model with the BonA-27N dimer structure modeled consistent with the observed decameric oligomerization (bottom). (G) A bead model of the BonA-27N monomer modeled from SAXS data (top), and a mesh representation of this bead model with monomer BonA-27N structure modeled. (H) The SAXS scattering curves for full-length BonA (top) and BonA-27N (bottom) in black, with simulated scattering curves for the bead models in panel A and B shown in green. (I) Class averages generated from negative stain EM images of the crosslinked BonA decamer showing a pentameric organization; Scale bar = 100 Å.

To understand the basis of oligomerization of BonA, both full-length and BonA-27N were analyzed via size-exclusion coupled small-angle X-ray scattering (SEC-SAXS) (Figure S5, Table S6). Despite the C-terminal extension, which largely lacks predicted secondary structure and was disordered in the BonA-27N crystal structure, SAXS scattering indicates that decameric BonA forms a compact particle in solution with maximum dimensions of ∼164 Å (Figure S5C, D). In contrast, SAXS scattering indicates that BonA-27N is highly flexible in solution with maximum dimensions of 107 Å, which is indicative of an unstructured and fully extended C-terminus (Figure S5G, H). These differences between decameric and monomeric BonAsuggest that intermolecular interactions stabilize the C-terminus of the oligomeric form of the protein.

Molecular envelopes of full-length and BonA-27N were modeled based on SAXS scattering data. For full-length BonA, C5 symmetry was imposed, based on the decameric organization of the oligomer and the dimer observed in the crystal structure. The resulting molecular envelope was prolate, with dimensions of ∼172 by 102 Å. Five dimers of the BonA-27N crystal structure could be modeled with C5 symmetry into a bulge at the center for the envelope. The N and C-termini of all molecules are orientated in the same direction, which is required by the lipid anchored N-terminus of BonA. Regions truncated or disordered in the BonA-27N crystal structure could be accommodated by the remainder of the molecular envelope (Figure 4F). The molecular envelope of BonA-27N was indicative of a monomer, with dimensions of ∼133 by 40 Å. The crystal structure of BonA-27N could be modeled into a bulge at one end of the envelope, with additional space accounting for the unstructured C-terminal extension (Figure 4G). The simulated scattering curves for both envelopes were an excellent fit for the experimental data (Figure 4H).

To validate our SAXS based modeling of the BonA decamer, we further investigated full-length BonA via negative-stain electron microscopy (NS-EM). Initial analysis of EM-grids prepared with native BonA did not contain discrete particles. To stabilize the decamer, on-column glutaraldehyde crosslinking was performed, stabilizing BonA as first a dimeric and then a decameric species with increasing glutaraldehyde concentration (Figure S4B). NS-EM of the crosslinked sample revealed largely uniform monodisperse particles (Figure S4C). 2D-class averages derived from these images are suggestive of a particle with dimensions compatible with the BonA SAXS envelope and C5 symmetry as predicted by other analyses (Figure 4I).

## Discussion

In this work, we identify BonA, a novel member of the bacterial dual-BON domain family of proteins, produced by *A. baumannii* and encoded by other members of the family *Moraxellaceae*. We demonstrate that BonA is anchored to the outer membrane where it plays a role in maintaining membrane structure and is required for twitching motility. Through structural analysis, we show that BonA possesses unique structural features and forms a divisome localized decamer that likely mediates its function (Figure 5). We show that while BonA shares a common outer membrane and divisome localization to DolP from *E. coli* and *Neisseria* spp. [26, 29, 35], its loss does not lead to the gross defects in outer membrane permeability observed in DolP deletion mutants. Furthermore, while DolP is monomeric and mediates its function and localization via phospholipid binding [28], BonA is a decamer that lacks the conserved lipid-binding residues found in DolP [28]. These differences suggest a role for BonA in outer-envelope function that is distinct from that of DolP and indicates functional divergence within the dual-BON domain protein family.

**Figure 5.**
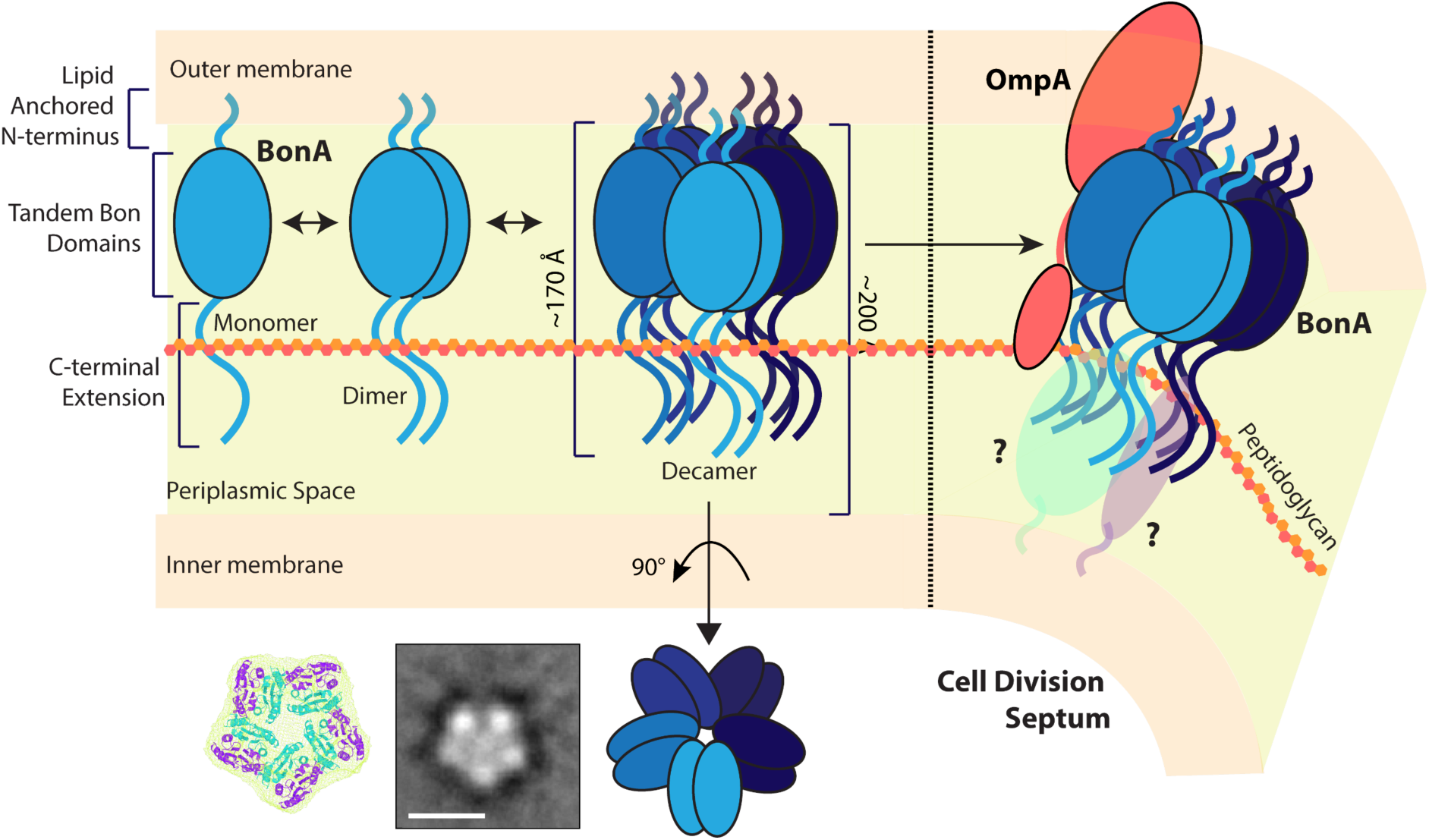
A model of BonA localization, oligomerization, and potential function at the outer membrane. BonA is anchored to the periplasmic side of the outer membrane where it forms a transient decamer that spans the majority of the periplasmic space. BonA is recruited to the site of cell division where it may interact with the peptidoglycan and act as a membrane-spanning scaffold for divisome proteins.

The change in outer membrane density associated with the loss of BonA suggests a significant alteration in the structure or composition of this membrane or the physical membrane-peptidoglycan links. Consistent with this, the loss of twitching motility observed in *A. baumannii* ATCC 17978 *ΔbonA*, likely mediated by an outer-envelope spanning type IV pilus [44], is suggestive of a perturbed outer envelope. These data are also consistent with our previous finding that BonA is upregulated in response to outer membrane destabilizing polymyxins [40, 41], and broadly indicates a role for BonA in supporting optimal outer membrane function.

While further work is required to determine the precise role of BonA in outer membrane function, our structural analysis provides important insights into BonA function. We show that BonA forms a decamer that is ∼172 Å in length. In the context of the periplasmic space, where the nominal distance between the outer and inner membranes is ∼200 Å [48], outer membrane-anchored BonA would span the majority of the periplasm if extending perpendicular from the membrane (Figure 5). In this configuration, BonA would penetrate the peptidoglycan layer and would be capable of interacting with proteins embedded in the inner membrane, thus bridging the inner and outer membranes. When localized to the site of cell division, BonA could tether the outer membrane to the peptidoglycan or the membrane-spanning divisome complex (Figure 5). In support of this hypothesis, in-cell crosslinking data shows interactions occur between BonA and OmpA in *A. baumannii* [47], with OmpA playing a role in tethering the outer membrane to the peptidoglycan [49]. The transient nature of BonA oligomerization is also consistent with a role in coordinating a dynamic process during cell division. If BonA is indeed important for coordinating the outer envelope during cell division, its loss would lead to improper remodeling of this structure, which is consistent with the *ΔbonA* phenotypes we observe.

The cell envelope provides a key defense for *A. baumannii* against antimicrobial compounds and environmental stress. To effectively combat *A. baumannii* infection and its persistence in the hospital environment, we must develop strategies to overcome the outer envelope’s defenses. To do so, a robust understanding of the key factors required for outer membrane construction and maintenance is required. Our work on BonA informs this understanding and provides insights into the unique role of this protein in supporting outer membrane function in *A. baumannii*.

## Materials and Methods

### Protein sequence analysis

To determine the relationship between distantly related dual-BON domain family members we constructed a tree of BonA homologs, identified with a HMMER search of the reference proteomes database using BonA as a query sequence [43]. BonA homologs identified in the HMMER search were curated to only include proteins with a dual-BON domain architecture and a lipobox sequence determined using SignalP 5.0 [50]. This yielded 896 sequences, which were further reduced for tree construction using CD-Hit to filter sequence with a pairwise similarity of <75 %, yielding 565 sequences (Table S1) [51]. These 565 protein sequences, plus OsmY from *E. coli* as a sequence to define the root branch, were aligned using MUSCLE [52] implemented in the phylogenetic analysis program MEGAX (v.10.1.7) [53], which was subsequently used as the input for constructing a maximum likelihood (ML) phylogenetic tree to infer evolutionary relationships for this protein family. The best amino acid substitution model was inferred using MEGAX which compared 56 different models; the Jones-Taylor-Thornton (JTT) model with a gamma distribution of 5 discrete gamma categories and invariant sites (G + I) was selected. To infer tree topology, the default ML heuristic method ML Nearest-Neighbor-Interchange (NNI) was applied and initial trees were made with Neighbour-Joining and BioNJ algorithms. The final tree was built by including all residues and bootstrapping with 100 replicates.

### Strain propagation, maintenance, and antimicrobial susceptibility testing

*E. coli* and *A. baumannii* were propagated in lysogeny broth (LB) and LB agar at 37°C. Antimicrobial susceptibility was conducted per CLSI guidelines using the broth microdilution method and cation adjusted Muller Hinton broth. Minimum inhibitory concentrations (MIC) were defined as >80% reduction in growth, and significance considered as >2 concentration increase or decrease in MIC relative to the wild type control.

### Construction of *ΔbonA* strains in *A. baumannii*

Plasmid DNA, genomic DNA, and PCR products were purified using relevant kits from Bioneer, QIAGEN, and Promega, respectively, following the manufacturer’s instructions. The *A. baumannii ΔbonA* mutants were constructed as described previously [54], with minor modifications. Briefly, the kanamycin resistance cassette was PCR amplified from pKD4 using disruption primers containing >80 bases of homology to the *bonA* flanking sequence (as described in Table S6). The resultant fragments were gel purified and introduced into *A*.*baumannii* strains ATCC17978 and ATCC19606 by electroporation as previously described [55], with selection on LB agar supplemented with 50 μg/ml kanamycin. The mutations were confirmed by PCR amplification using primers flanking the insertion, followed by Southern hybridization of genomic DNA digested with *Eco*RV, probed with kanamycin and *bonA* specific DIG-labeled probes, as described previously [56].

For complementation, the full-length *bonA* sequence plus 500 nucleotides upstream of the translational start site (deemed to include the native promoter) were PCR amplified from ATCC19606 with forward and reverse complementation primers encoding 5’ *Aat*II and *Eco*RI restriction sites, respectively. The resultant fragments were digested and ligated into the *E. coli-Acinetobacter* shuttle vector, pWH1266 [57]. The p*BonA* constructs were confirmed by sequencing before electroporation into the respective mutant strains as described previously, with the pWH1266 vector-only used as a control.

### Twitching motility assays

Twitching motility was assessed as described previously [44]. Briefly, a 1 μl drop of stationary phase culture was placed onto the center of a 0.25% modified LB agarose and incubated at 37°C for up to 48 hours. Three independent experiments were performed for each.

## BonA antisera generation

Polyclonal rabbit antisera for the detection of BonA was generated at the Monash Animal Research Platform, from recombinant proteins purified in-house. Rabbits were serially injected with purified protein (10 mg/ml) in combination with complete (first injection) or incomplete (subsequent injections) Freund’s adjuvant, over 1-3 months, with clarified rabbit sera periodically tested for reactivity to the target protein. Once acceptable levels of reactivity were achieved rabbits were euthanized and clarified sera were collected and stored at -80 °C.

### Isolation and fractionation of membranes from *A. baumannii*

*A. baumannii* cells were cultured in LB media and grown to an OD_600_ of ∼0.6 before harvesting. Membranes were purified and subsequently fractionated by sucrose density fractionation (60:55:50:45:40:35% w/w) as described previously [58].

### Detection and localization of BonA in *A. baumannii* cell extracts via Western Blot

For the detection of BonA in cell extracts, 50 µg of isolated total membranes were analyzed by 10% SDS-PAGE or 5-16% blue-native (BN)-PAGE [59] and was subsequently analyzed by Western blotting against BonA (antibody dilution - 1:20,000). To determine the cellular localization of BonA, 30 µl aliquots of each fraction from the sucrose gradient were separated by 10% SDS-PAGE for Coomassie staining and subsequent Western blotting as described above.

### Localization of BonA in *A. baumannii* cells by immunofluorescence microscopy

Bacterial cultures were grown to mid-log phase in LB media at 37°C with shaking (200 rpm). Then 500 μl of culture media was centrifuged (4,000 × g, 5 min, 4°C), washed twice in PBS, and resuspended in 500 μl of PBS. 8-well, coverglass-bottom chambers (Sarstedt) were coated with 0.01% (v/v) poly-L-lysine (Sigma-Aldrich, P8920) for 10 min at room temperature before excess poly-L-lysine was removed. Afterward, 200 μl of bacterial cell suspension was immobilized onto each well. To ensure a monolayer of bacteria was formed at the bottom of each well, chamber slides were subjected to centrifugation (4,000 × g, 3 min, 4°C), followed by several washing steps to remove non-adhered cells. The monolayer of bacteria was then fixed with a mixture of paraformaldehyde (2% w/v) and glutaraldehyde (0.2% v/v) in PBS for 5 min at 4°C, which was then washed with PBS to remove excess fixatives. To reduce auto-fluorescence caused by the background, samples were treated with a fluorescence quencher, 0.1% (w/v) NaBH_4_ in PBS for 15 min, followed by several washing steps of PBS. Samples were then permeabilized with Triton X-100 (0.001% v/v in PBS), followed by three washing steps with PBS.

Before antibody staining, samples were blocked with 5% (w/v) BSA in PBS for 1 hr at room temperature, followed by incubation with anti-BonA antisera (1:1,000 in 5% w/v BSA in PBS) for 1-hr mixing by rotary inversion at room temperature. Samples were washed thoroughly with PBS to remove excess antiserum. Secondary staining was carried out for 45 mins at room temperature using anti-rabbit immunoglobulin G (IgG)-Alexa Fluor 488 (ThermoFisher®, A-11008) diluted to 1:3,000 (in 5% BSA in PBS), followed by several washing steps to remove excess antibody. Olympus IX-81 inverted fluorescence microscope equipped with Olympus Cell^M software was used to visualize bacteria samples using the 100× objective with fluorescein isothiocyanate (FITC) filter.

### Protein Expression and Purification

DNA encoding full-length BonA and Bon*A*-C45 were amplified by PCR, with C-terminal NcoI and XhoI restriction sites and cloned into a pET20b derived vector which added a 10x N-terminal His-tag followed by a TEV cleavage site, via restriction digest and ligation. The resulting vector was transformed into *E. coli* BL21 (DE3) C41 cells. DNA encoding BonA-27N, and BonA-27N-45C were amplified by PCR, minus stop codon, with C-terminal NdeI and XhoI restriction sites and cloned into pET22b vector which added a 6x C-terminal Histag. The resulting vectors were transformed into *E. coli* BL21 (DE3) C41 cells. Protein expression was performed in terrific broth (12 g tryptone, 24 g yeast extract, 61.3 g K_2_HPO_4_, 11.55 g KH_2_PO_4_, 10 g glycerol) with 100 mg/ml ampicillin for selection. Cells were grown at 37°C until OD_600_ of 1.0 induced with 0.3 mM IPTG and growth for a further 14 hours at 25°C. For selenomethionine labeled BonA-27N, the BonA-27N construct was transformed into the methionine auxotrophic *E. coli* strain Crystal Express (DE3). Cells were grown in M9 minimal media containing 100 mg/l of each amino acids (minus methionine) and 50 mg/l selenomethionine. Cells were harvested by centrifugation, lysed using a cell disruptor (Emulsiflex) in Ni-binding buffer (50 mM Tris, 500 mM NaCl, 20 mM imidazole [pH 7.9]) plus 0.1 mg/ml lysozyme, 0.05 mg/ml DNAse I, and cOmplete protease cocktail inhibitor tablets (Roche). The resulting lysate was clarified by centrifugation and applied to Ni-agarose resin, followed by washing with 10x column volumes of Ni-binding buffer, and elution of bound proteins with a step gradient of Ni-gradient buffer (50 mM Tris, 500 mM NaCl, 500 mM Imidazole [pH7.9]) of 5, 10, 25 and 50%. Eluted fractions containing recombinant protein were pooled and applied to a 26/600 Superdex S200 size exclusion column equilibrated in SEC buffer (50 mM Tris, 200 mM NaCl [pH 7.9]). The recombinant protein was then pooled concentrated to 10 mg/ml, snap-frozen, and stored at -80 °C.

### Size-exclusion chromatography multiangle light scattering (SEC-MALS)

The absolute molecular masses of BonA-FL and truncated variants were determined by SEC-MALS. 100-μl protein samples (1-5 mg/ml) were injected onto a Superdex 200 10/300 GL size-exclusion chromatography column in 20 mM Tris, 200 mM NaCl [pH 7.9] at 0.6 ml/min with a Shimadzu LC-20A. The column eluent was fed into a DAWN HELEOS II MALS detector (Wyatt Technology) followed by an Optilab T-rEX differential refractometer (Wyatt Technology). Light scattering and differential refractive index data were collected and analyzed with ASTRA 6 software (Wyatt Technology). Molecular masses and estimated errors were calculated across individual eluted peaks by extrapolation from Zimm plots with a dn/dc value of 0.1850 ml/g. SEC-MALS data are presented with absorbance (280 nm) plotted alongside fitted molecular masses (Mr).

### Protein crystallization, data collection, and structure solution

Purified BonA proteins were screened for crystallization conditions using commercially available screens (approximately 800 conditions). Crystals grew from drops containing BonA-27N in conditions containing 0.2 M Zn Acetate, 0.1 M Na Acetate, 20 % PEG 3350 [pH 4.5], crystals were optimized in this condition. Crystals were cryoprotected by increasing PEG 3350 concentration to 30% and flash cooled in liquid N_2_. Diffraction data were collected at 100 K at the Australian Synchrotron on selenomethionine labeled crystals and processed in the space group P3_1_21 to 1.65 Å. Heavy atom sites were located, phases were obtained using single-wavelength anomalous dispersion (SAD) and the initial model was built using Autosol from the Phenix package [60]. Eight heavy atom sites were located, 4 of these sites were Selenium, 4 of these sites were Zn. The BonA-N27 model was improved manually in Coot and refined using Phenix refine and Refmac [60-62]. Analysis of the BonA-27N crystal structure was performed using the Phenix and CCP4 packages, non-crystallographic interfaces were predicted using PISA [46, 60, 63].

### Small Angle X-ray Scattering

Small-angle X-ray scattering (SAXS) was performed using Coflow SEC-SAXS at the Australian Synchrotron [64]. Purified BonA and BonA-27N were analyzed at a pre-injection concentration of 10 mg/ml. Scattering was collected over a *q* range of 0.0 to 0.3 Å^-1^. A buffer blank for each SEC-SAXS run was prepared by averaging 10-20 frames pre- or post-protein elution. Scattering data from peaks corresponding to BonA and BonA-27N were then buffer subtracted and scaled across the elution peak, and compared for inter-particle effects. Identical curves (5-10) from elution were then averaged to provide curves for analysis. Data were analyzed using the PRIMUS package, ScÅtter, and DAMMIF modeler [65].

### Analytical ultracentrifugation

Sedimentation velocity (SV) was carried out in a Beckman Coulter Optima analytical ultracentrifuge using an An-50 Ti 8-hole rotor. BonA-FL (370 μl) at concentrations ranging from 0.25 to 2 mg/ml was loaded into a 12 mm path-length centerpiece and centrifuged at 40,000 rpm for ∼6 h at 20°C. Scans were collected every 20 seconds using absorbance optics (at 230, 240, and 280 nm; a radial range of 5.8 - 7.2 cm, and radial step-size of 0.005 cm). 50 mM Tris, 200 mM NaCl, pH 7.9 was used as the buffer. Data were analyzed with SEDFIT using the continuous c(s) distribution model [66]. SEDNTERP was used to calculate the partial specific volume, the buffer density, and viscosity at 15°C and 20°C.

### On column crosslinking and negative-stain electron microscopy

To stabilize the BonA decamer an ‘on-column’ crosslinking method was used. Initially, 200 µl of glutaraldehyde solution (0.05-0.5% in dH_2_O) was injected to a pre-equilibrated Superdex 200 10/300 column in buffer (20 mM HEPES, 150 mM NaCl, [pH 7.4]). The column was run at 0.25 ml/min for 20 min (5 ml buffer). Subsequently, the column flow was paused, and the injection loop was flushed using buffer followed by injection of purified BonA (200 µl, at 10 mg/ml). Subsequently, the column was run at 0.25 ml/ min and 0.5 ml fractions were collected. Collected fractions were immediately quenched by the addition of 50 µl of 50 mM Tris, pH 7.5. Crosslinking efficiency was visualized by running the individual fractions on a 10 % SDS gel and cross-linked fractions were flash-frozen for NS-EM analysis.

Native and crosslinked BonA were serially diluted in buffer (20 mM HEPES, 150 mM NaCl, pH [7.4]) and 5 µl was spotted onto freshly glow-discharged carbon-coated 200-mesh copper grids (PELCO), followed by blotting to remove all but a thin film of protein solution. Blotted grids were fixed with the tungsten-based Nano-W strain (Nanoprobes), by adding the stain to each grid, followed by 60 seconds incubation and blotting, repeated 3 times before air drying. The grids were imaged on a 120 keV Tecnai Spirit G2 microscope (FEI) equipped with a 4 K FEI Eagle camera. Images were processed, particles were picked and 2D classes generated using the RELION package (V 2.1) [67].

## Acknowledgments

This research was undertaken on the MX1, MX2, and SAXS/WAXS beamlines at the Australian Synchrotron, part of ANSTO (CAP12312, and M12480). We would like to thank the Monash Molecular Crystallisation Facility for their assistance with sample characterization, crystallographic screening, and optimization. We would like to thank Mr. Hari Venugopal and the Ramaciotti Centre for Cryo-Electron Microscopy for their assistance with electron microscopy experiments. We would like to thank Dr. Pankaj Deo for his assistance with fluorescence microscopy and Dr. Sarah Atkinson for her assistance with the AUC measurements.

## Data availability

The crystallographic coordinates and associated structure factors for BonA are available at the Protein Data Bank (PDB) with the accession code 6V4V. Small-angle X-ray scattering data for BonA full-length and BonA-27N are available in the SASBDB with the accession codes SASDJW3 and SASDJX3. Accession numbers of protein sequences used to construct the phylogenetic tree are provided in Table S1.

## Author Contributions

Conceived and designed the experiments: RG, FCM, RAD, EH, TL

Performed the experiments: RG, FCM, RAD, MB, DG

Analyzed the data: RG, FCM, RAD, MB, DG, PML, EH, TL

Contributed reagents/materials/analysis tools: RG, SB, HV, AYP, CG, JL, EH, TL

Wrote the paper: RG, TL

Edited and approved the manuscript: All authors

## Supporting Information Legends

### Supplemental Figures

**Figure S1.**
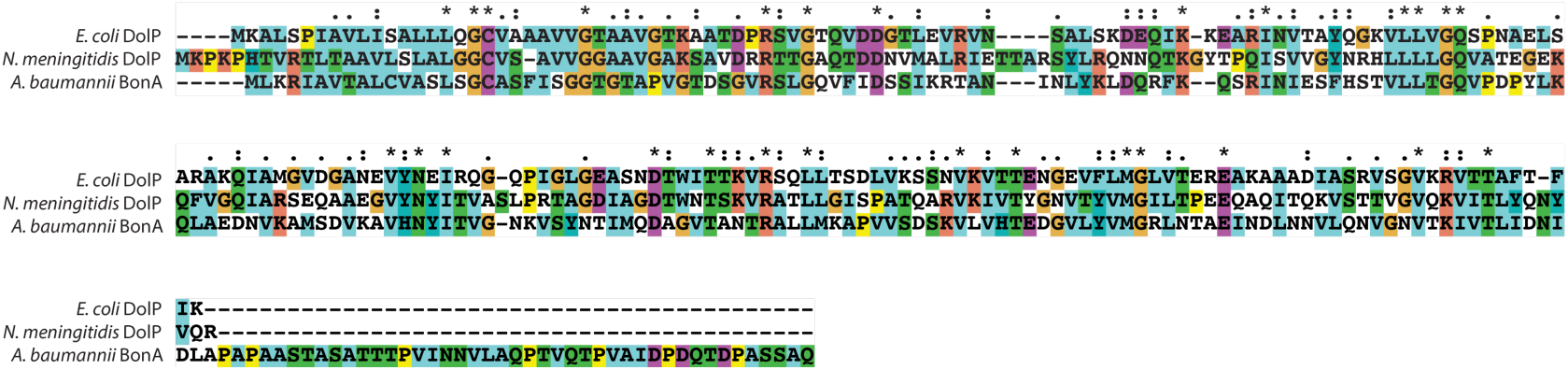
Multiple sequence alignment of BonA from *A. baumannii* and DolP from *E. coli* and *N. meningitidis*. The proline-rich C-terminal extension present in BonA but absent from the *E. coli* and *N. meningitidis* proteins is notable.

**Figure S2.**
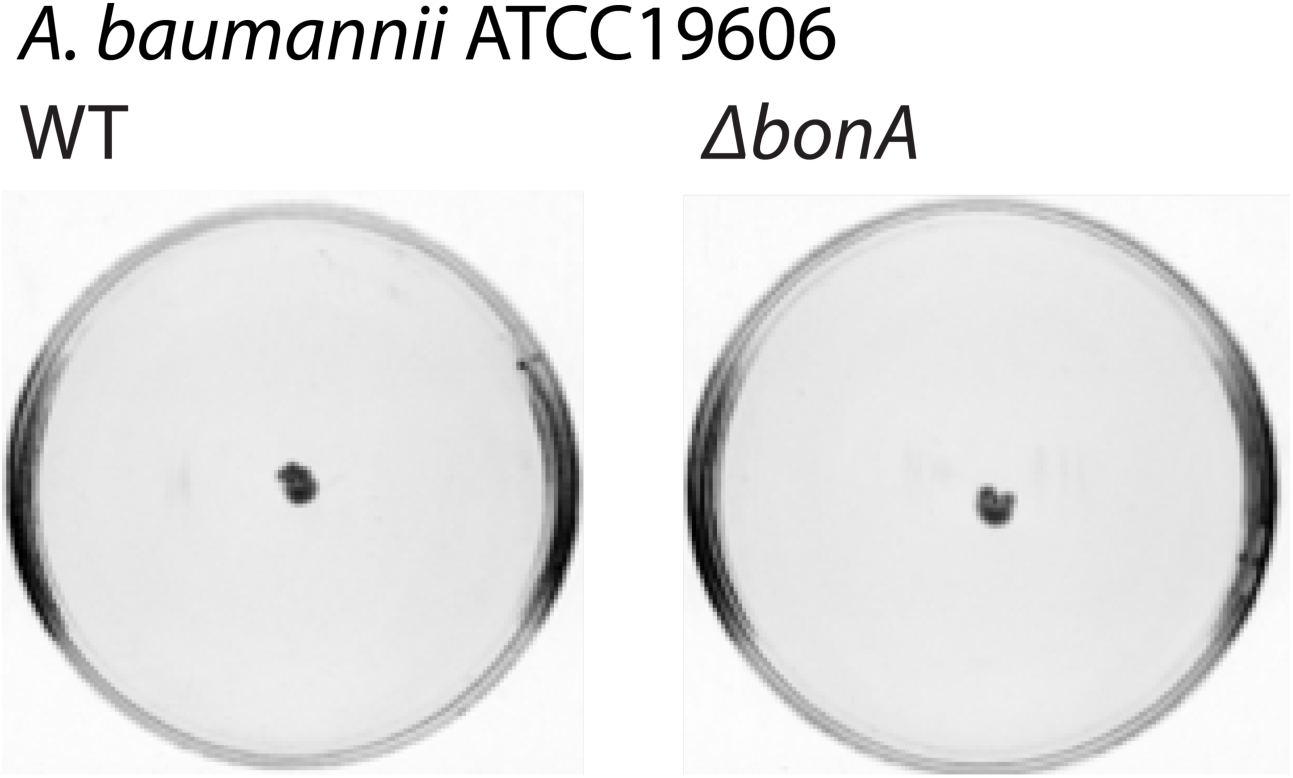
Semi-solid motility assay plates of *A. baumannii* ATCC19606. Showing that the wildtype strain is non-motile and therefore this phenotype is unaffected by the loss of BonA

**Figure S3.**
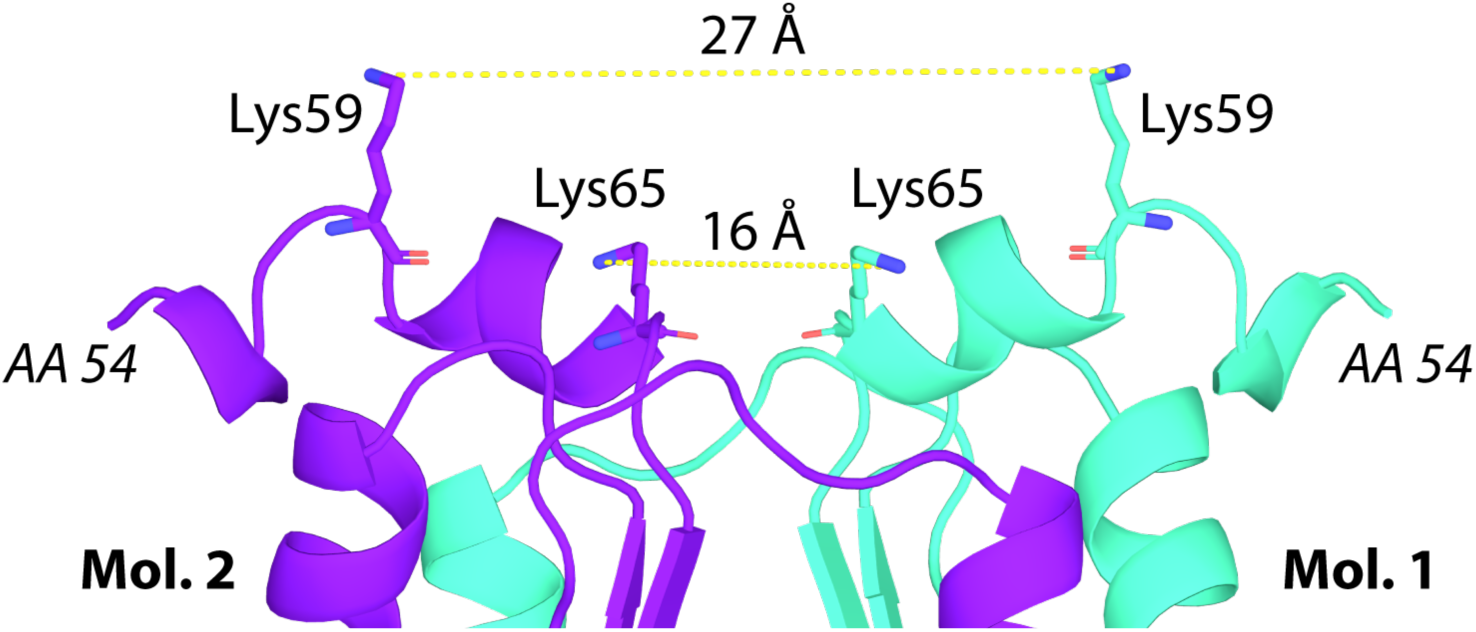
Lysine residues of BonA identified as interacting in vivo cross-linking conducted by Wu et. al. [47], shown on the structure of BonA-N27.

**Figure S4.**
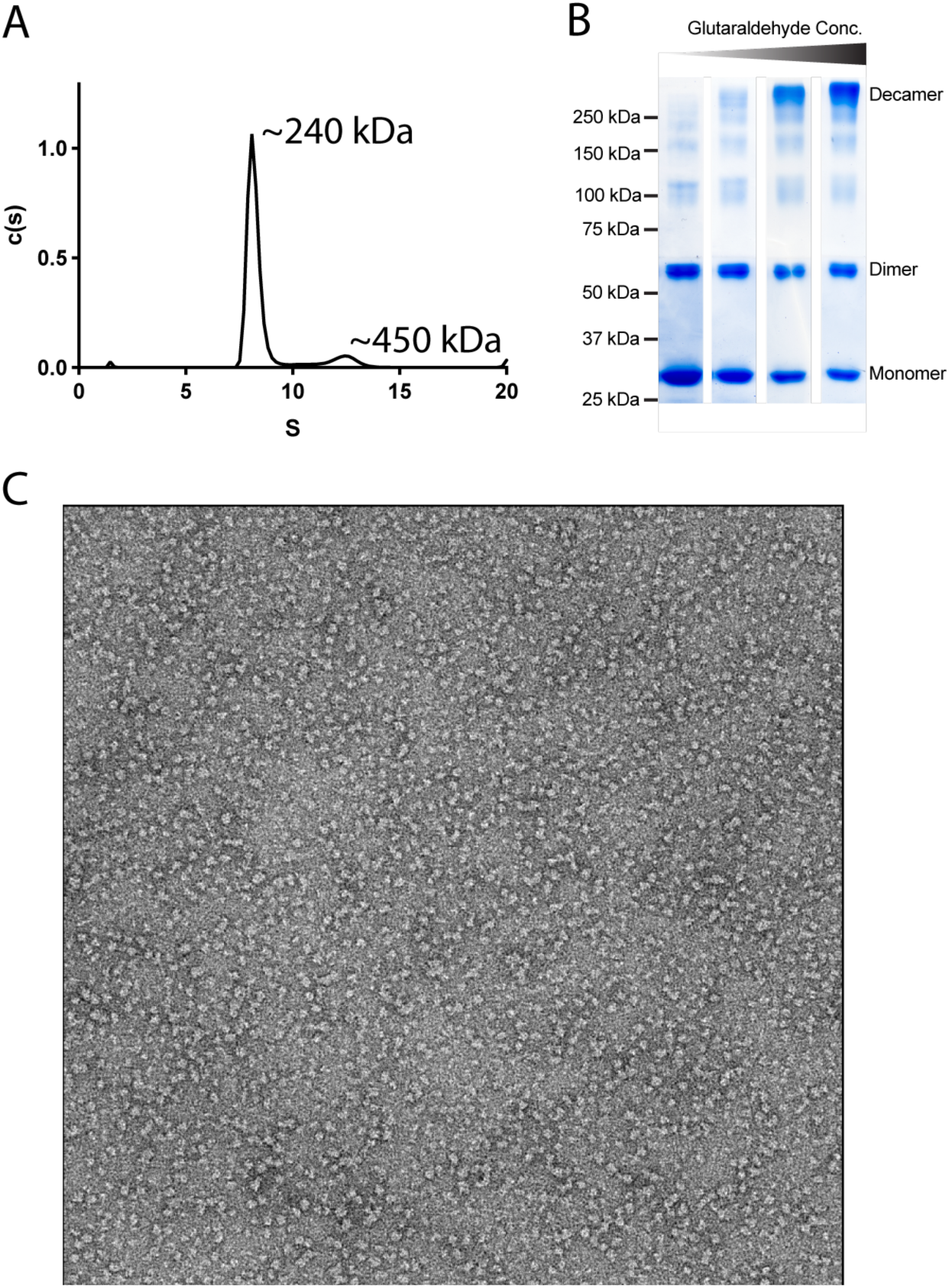
Full-length BonA forms a decameric oligomerization consisting of discrete compact particles. (A) Analytical ultracentrifugation sedimentation profile for full-length BonA showing that it exists predominantly as a 240 kDa species, consistent with decameric oligomerization. (B) An SDS-PAGE gel containing the eluted peak fraction from on column crosslinking of full-length BonA, with increasing concentration of the crosslinking reagent glutaraldehyde from left to right. (C) A representative negative stain EM image of crosslinked full-length BonA from the highest glutaraldehyde concentration (0.5 %) shown in panel B.

**Figure S5.**
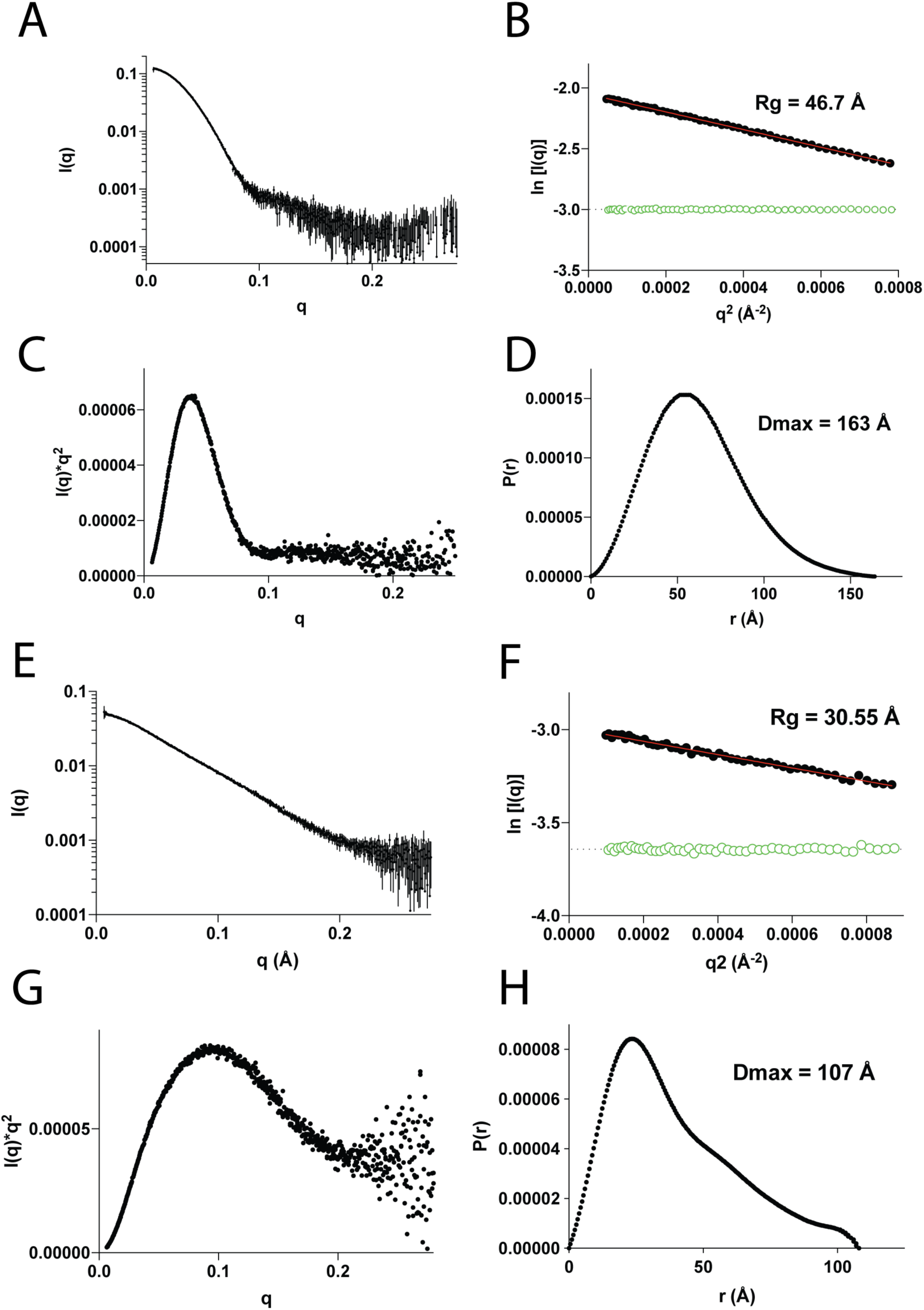
SAXS scattering data plots for full-length BonA and BonA-27N. The SAXS scattering curve for full-length BonA (A), and the Guinier (B), Kratky (C) and P(r) (D) plots derived from it. The SAXS scattering curve for BonA-27N (E), and the Guinier (F), Kratky (G), and P(r) (H) plot derived from it.

## Supplemental Tables

Table S1 Sequence identity matrix of dual-BON family proteins from *A. baumannii, E. coli* and *N. meningitidis*

Table S2 BonA homologs utilized for phylogenetic tree generation

Table S3 Susceptibility of *Δ*bonA *A. baumannii* strains to selected antimicrobial agents

Table S4 Crystallographic data collection and refinement statistics

Table S5 BonA-27N dimer interface statistics calculated by PISA[46]

Table S6 Small-angle X-ray scattering data collection and modeling statistics

Table S7 Primers and strains utilized for this study

